# Caffeinated Coli: Inspiring the Next Generation of Scientists through Synthetic Biology

**DOI:** 10.64898/2026.07.17.738993

**Authors:** Neil Yao Tian, Riya Reddy, Connie Yu Zhang, Bailey Williams, Dennis Michael Mishler

## Abstract

The Caffeinated Coli Educational Module brings inquiry-driven learning to high schools, introducing students to scientific research, synthetic biology, and genetic engineering. The module focuses on the exploration of a genetically engineered strain of *E. coli* modified to grow exclusively on caffeine and, as such, can be used as a measurement device to determine the amount of caffeine in a liquid or beverage. Students conduct two bioassay experiments using these bacteria. During this process, they learn to create cultures and then measure bacterial growth followed by calculating the caffeine concentrations of unknown samples using their own data. This flexible module contains five weeks of original lectures and student learning materials that follow Next Generation Science Standards (NGSS), allowing the content to be adapted for basic, intermediate, or advanced biology courses. Upon completion of the module, students and teachers expressed that the most memorable aspects of the module include collaboration with peers, hands-on learning of content, and the opportunity to interact with the professor/mentors through office hours. Already implemented in 8 Texas high schools over the past two academic years, our module inspires STEM learning while bringing 21^st^ century biology research to new audiences.

---

*Jurassic Park’s* cloning of dinosaurs. Spider-goats’ production of spider silk (Jones, 2010). Impossible Food’s plant-based burgers (Meat-free Outsells Beef, 2019). Only one of these is science fiction. The others are remarkable examples of what synthetic biology can achieve. Synthetic biology is a field of science where researchers genetically modify organisms to perform new functions or to have new properties that were once just a figment of our imagination. Although a rapidly progressing branch of science, most students in the United States are not introduced to synthetic biology until college, if ever. Unlike robotics programs or math clubs, which are commonplace in junior highs and high schools, synthetic biology has little to no footing in these early educational environments, and even that is primarily through specialized summer programs or international competitions (Menard, 2024).

## Caffeinated Coli

The Caffeinated Coli Module builds on work by Quandt and colleagues, which was itself inspired by an undergraduate research project called “Caffeinated Coli” (Quandt et al., 2013). This work created a biosensor in the form of a modified strain of *Escherichia coli* capable of measuring caffeine and related methylxanthines, demonstrating a practical application of synthetic biology. The engineered strain lacks the *guaB* gene, which is required for the *de novo* synthesis of guanine, an essential nucleic acid for DNA replication and cell growth. However, this strain (known as Δ*guaB E. coli*) can bypass this deficiency by converting xanthine into guanine using a native xanthine salvage pathway if supplemented with xanthine (Figure 1).

**Figure 1.**
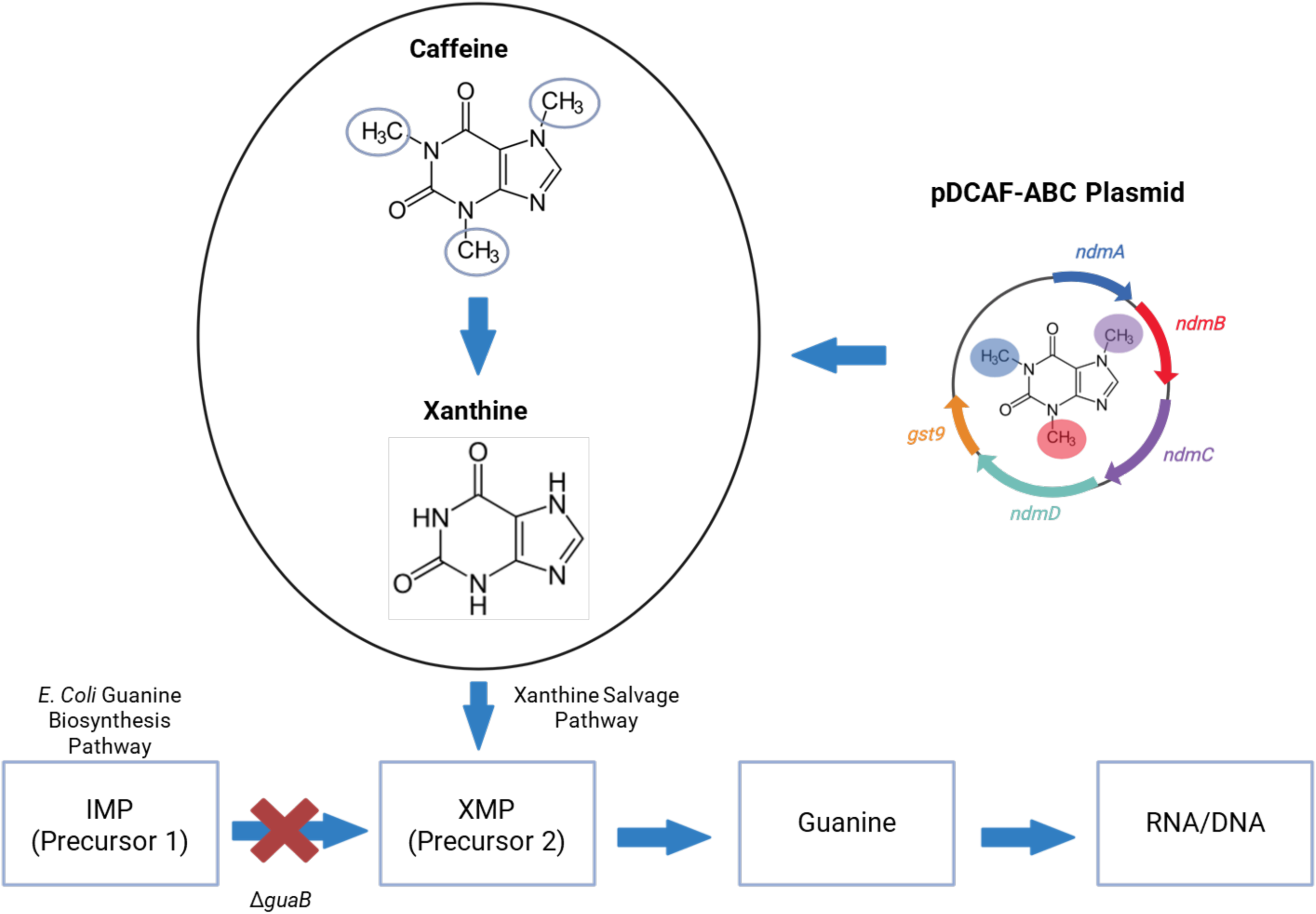
Novel Caffeinated Coli Guanine Biosynthesis Pathway.

To give *E. coli* the ability to metabolize methylxanthines into xanthine, researchers developed the pDCAF-ABC plasmid. This plasmid contains genes that encode three demethylase enzymes (NdmA, NdmB, and NdmC) capable of removing methyl groups from caffeine (1,3,7-trimethylxanthine), converting it to xanthine. The plasmid also includes the *ndmD* and *gst9* genes, which support this demethylation process (Quandt et al, 2013). When this plasmid is inserted into Δ*guaB E. coli*, the resulting bacteria are known as “Δ*guaB* + pDCAF-ABC *E. coli,*” commonly shortened to “pDCAF-ABC *E. coli*,” or more simply, “Caffeinated Coli”. Caffeinated Coli can metabolize methylxanthines, converting them to xanthine (Figure 1, top). This xanthine is ultimately turned into guanine sustaining growth in minimal media. Thus, since this strain of *E. coli* only grows when fed caffeine or another methylxanthine, it can be used as an accurate biosensor, on par with HPLC, to determine the amount of caffeine in a beverage (Quandt et al, 2013; Gutierrez et al, 2019). The pDCAF-ABC strain can be readily obtained from Addgene (Plasmid #113652), a non-profit, for a small distribution fee. Growth conditions are detailed at this site and in our educational materials, which include the experimental procedures as well as the teacher preparation guidelines.

Here, we present details of the 5-week educational module, which highlights the critical pieces of classroom instruction, including lesson plans, student activities, and wet-lab work. This project-based synthetic biology curriculum for high schools meets Next Generation Science Standards (NSTA, 2016). Additionally, the manuscript includes a summary of student/teacher feedback, which showed an increased interest in STEM research and careers. Ultimately, the goal of this work is to offer high school students an opportunity to engage in synthetic biology research through hands-on, inquiry-based learning, modeled after courses designed for college freshmen (Rodenbusch et al, 2016).

## The 5-week Module

The Caffeinated Coli Educational Module is a flexible 5-week module designed to introduce high school students to research, genetic engineering, and synthetic biology (Figure 2). The lessons are designed for students to build proficiency in four NGSS Performance Expectations: *HS-LS1-1, HS-LS1-6: From Molecules to Organisms: Structures and Processes, HS-LS2-2: Ecosystems: Interactions, Energy, and Dynamics, and HS-LS3-1: Heredity: Inheritance and Variation of Traits* ( Table 1; NGSS Lead States 2013). In addition to NGSS alignment, each lesson was originally designed with TEKS (Texas Education Agency, 2020) standards in mind to meet student learning benchmarks in Texas (Supplemental Table 1). Since synthetic biology requires discipline learning in both engineering and biology, Science and Engineering Practices (SEPs) are layered throughout the module (Supplemental Table 2; NGSS Lead States 2013). In the first two weeks, students participate in lectures, group discussions, and exercises to build a foundational understanding of biology and research concepts on topics including the Scientific Method, the Central Dogma, genetic engineering, and experimental design. They also learn about the Caffeinated Coli biosynthesis pathway and practice essential lab techniques, such as micropipetting and making dilutions.

**Figure 2.**
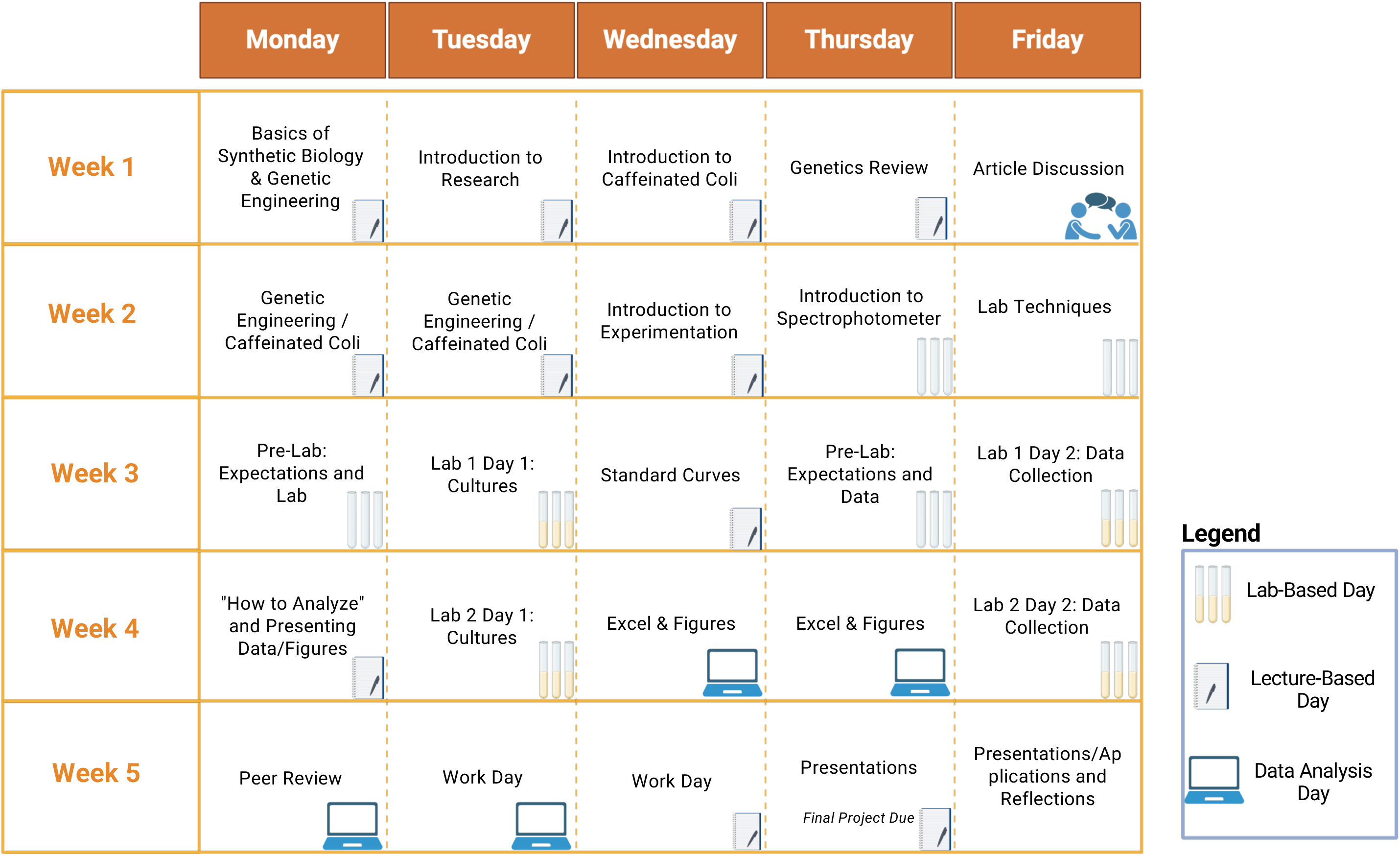
Overview of 5 Week Module.

**Table 1.** Lesson Alignment with NGSS Standards.

| Lesson | Next Generation Science Standards (NGSS) |
| --- | --- |
| **Week 1: Basics of Synthetic Biology** | HS-LS1-1. Construct an explanation based on evidence for how the structure of DNA determines the structure of proteins, which carry out the essential functions of life through systems of specialized cells. |
| **Week 1: Introduction to Caffeinated Coli** | HS-LS3-1: Ask questions to clarify relationships about the role of DNA and chromosomes in coding the instructions for characteristic traits passed from parents to offspring. |
| **Week 2: Introduction to ChromeBox** | HS-LS2-2. Use mathematical representations to support and revise explanations based on evidence about factors affecting biodiversity and populations in ecosystems of different scales. |
| **Week 3: Standard Curves** | HS-LS2-2. Use mathematical representations to support and revise explanations based on evidence about factors affecting biodiversity and populations in ecosystems of different scales. |

Students finish the second week by constructing the ChromeBox, an alternative device to a spectrophotometer, that they can use to measure bacterial growth. During Weeks 3 and 4, students apply their knowledge by conducting a bioassay to measure caffeine levels in beverages using genetically modified bacteria. Bacterial growth is then measured using either a spectrophotometer or their ChromeBox. Students then analyze their data using a spreadsheet and template that we provide. During the final week, students can peer-review each group’s results and present their findings as part of the module’s conclusion. The module is designed with 3 primary learning objectives: for students to explore fundamental research and synthetic biology concepts, learn basic lab techniques, and conduct data analysis to determine the amount of caffeine in a beverage (Figure 3).

**Figure 3.**
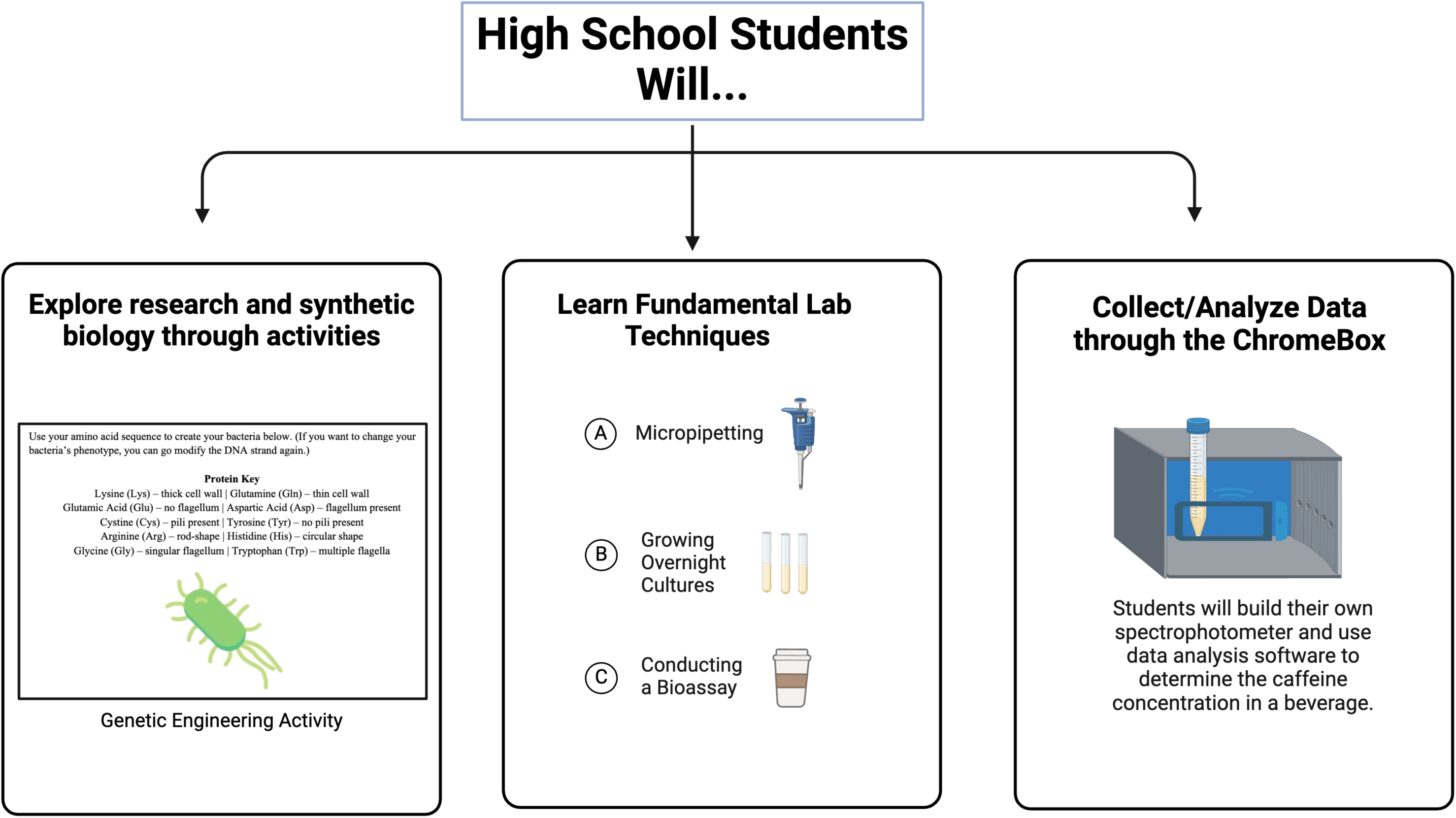
Learning Objectives for High School Students.

## Classroom Content (Weeks 1 and 2)

In Week 1 of the Caffeinated Coli Module, students build a foundational understanding of key concepts that will support the rest of the module. Week 2 delves deeper into the genetic engineering behind the creation of the Δ*guaB* + pDCAF-ABC strain of *E. coli* and introduces the lab techniques essential for the upcoming bioassay experiment. Lecture slides as well as student activities or readings for each day can be found online on our website for free (see Online Resources section).

Each day includes a lesson plan, a PowerPoint presentation, and a student learning activity (Table 2). Students begin each lesson with an initiating warm-up activity to expose them to the content and conclude with an exit ticket to gauge understanding. Each lesson is designed around the 5E lesson plan, which breaks down a lesson into five learning stages: engage, explore, explain, elaborate, and evaluate. The 5E lesson plans also include differentiation strategies for teachers to utilize if needed. A variety of activities are included to supplement students’ learning of advanced synthetic biology concepts, including example videos, group discussion activities, student learning activities, and exit ticket assessments to check for understanding. For instance, in the “Basics of Synthetic Biology” lesson (Table 3), students are introduced to general concepts with an engaging video detailing the hypothetical genetic engineering of Jurassic Park dinosaurs. Students will then explore the reality of synthetic biology’s capabilities as they discuss modern examples showcased in a PowerPoint presentation. Students solidify concepts regarding synthetic biology with an activity that explores the real attempts to revive the Woolly Mammoth (Callaway, 2024) using other living organisms’ DNA (Figure 4). Finally, students will answer a few free response questions about the various synthetic biology techniques and real-world uses that they learned that day. During the first weeks, students are also introduced to basic research methods, the mechanisms that underpin the Caffeinated Coli bioassay, the ChromeBox, and the importance of standard curves (Table 3).

**Table 2.**
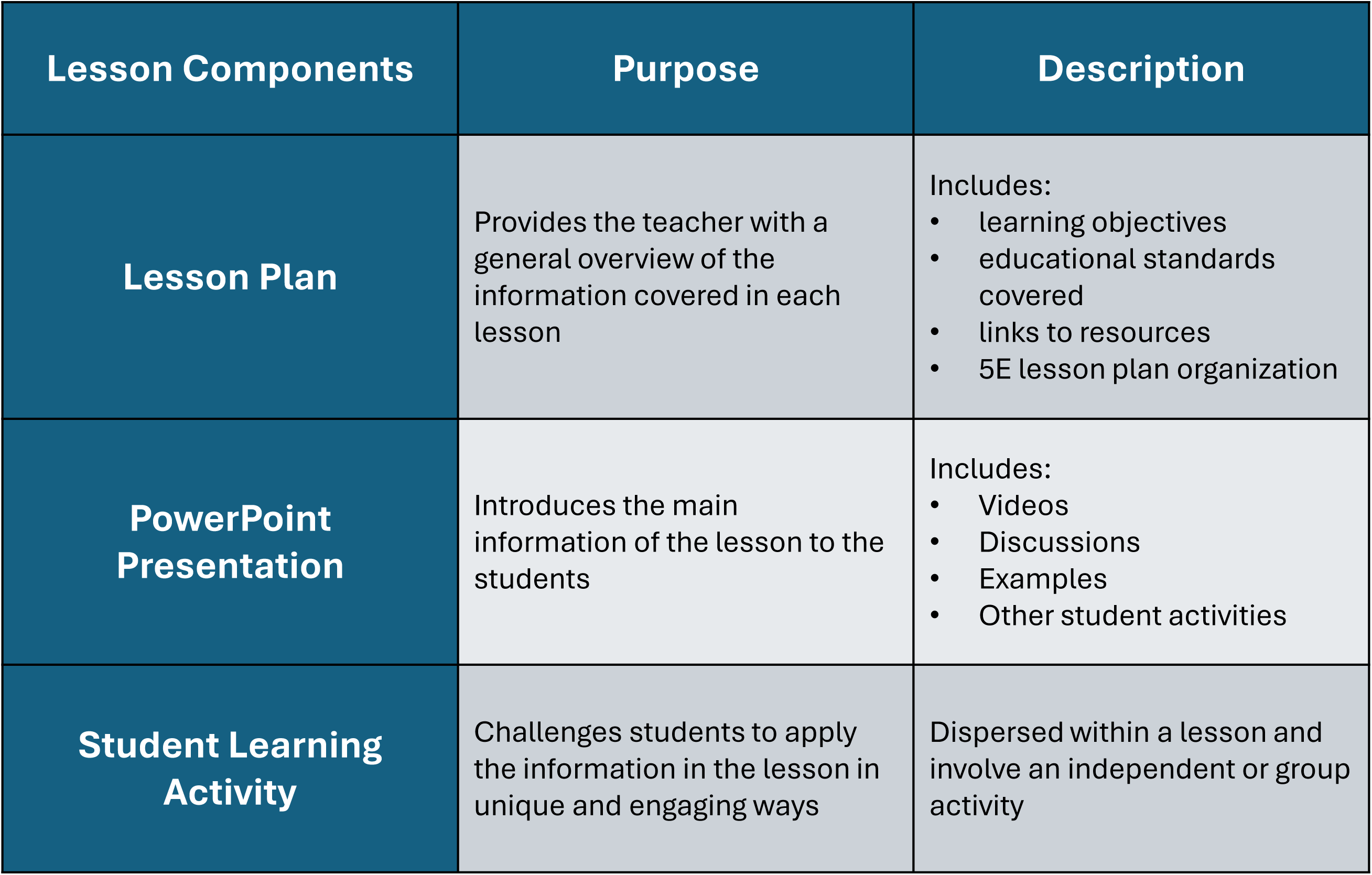
Components of Each Lesson: Lesson Plan, PowerPoint Presentation, and Student Learning Activity.

**Table 3.** Week 1, 2, and 3 Lesson Examples.

| Lesson | Lesson Plan Objective | PowerPoint | Student Activity |
| --- | --- | --- | --- |
| **Week 1: Basics of Synthetic Biology** | Introduces the field of Synthetic Biology through a guided discussion about Jurassic Park. | Includes a short Jurassic Park clip to introduce synthetic biology and examples of other synthetic biology projects including Dolly the Sheep, GloFish, and Spider Goats. | Reviving the Woolly Mammoth: students will follow the same synthetic biology premises behind Jurassic Park scientists' revival of dinosaurs to bring back the Woolly Mammoth. |
| **Week 1: Introduction to Caffeinated Coli** | Introduces the origins of Caffeinated Coli and the practical applications of this genetically modified strain of *E. coli*. | Poses questions for discussion about Caffeinated Coli and introduces background information on the topic. | Choose Your Drink: students will choose a beverage to measure the caffeine concentration of using Caffeinated Coli. |
| **Week 2: Introduction to ChromeBox** | Introduces students to the spectrophotometer and the alternative method of taking absorbance readings via the ChromeBox. | Starts with a guided discussion analyzing cultures with different bacterial growth and explains the mechanism behind the Spectrophotometer/ChromeBox. | Building the ChromeBox: students will create their own ChromeBox and identify the key structures of the ChromeBox. |
| **Week 3: Standard Curves** | Introduces students to standard curves by generating and analyzing their own data plots with classmates. | Begins with a warm-up activity analyzing examples of standard curves and introduces students to the Caffeinated Coli standard curve by showing its usefulness in predicting relationships. | Standard Curve Activity: students will measure each other's height and foot length and plot the data on a graph to observe potential relationships. |

**Figure 4.**
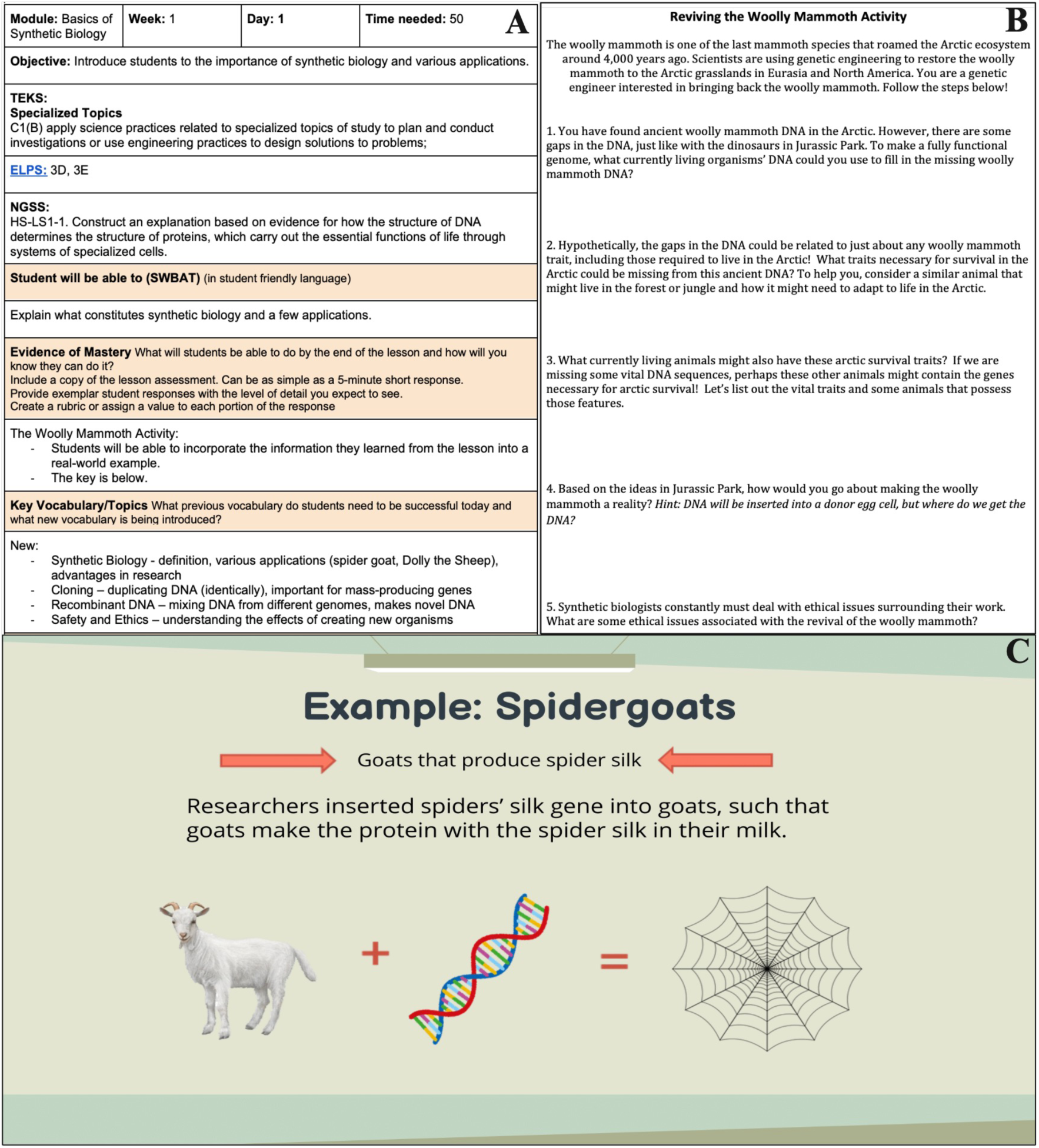
Basics of Synthetic Biology Curriculum Materials. Panel A: Lesson Plan; Panel B: Student Learning Activity; Panel C: PowerPoint Example.

## Caffeinated Coli Bioassay (Weeks 3 and 4)

In weeks 3 and 4, students move into the wet lab to conduct a bioassay, using the knowledge from previous weeks to measure caffeine concentration in a beverage. Conducting the bioassay requires several materials, including micropipettes/tips, dropper pipettes, genetically engineered *E. coli*, caffeine, pre-made plates, minimal media, caffeine/calibration stocks, plastic culture tubes, and glove boxes (Figure 5).

**Figure 5.**
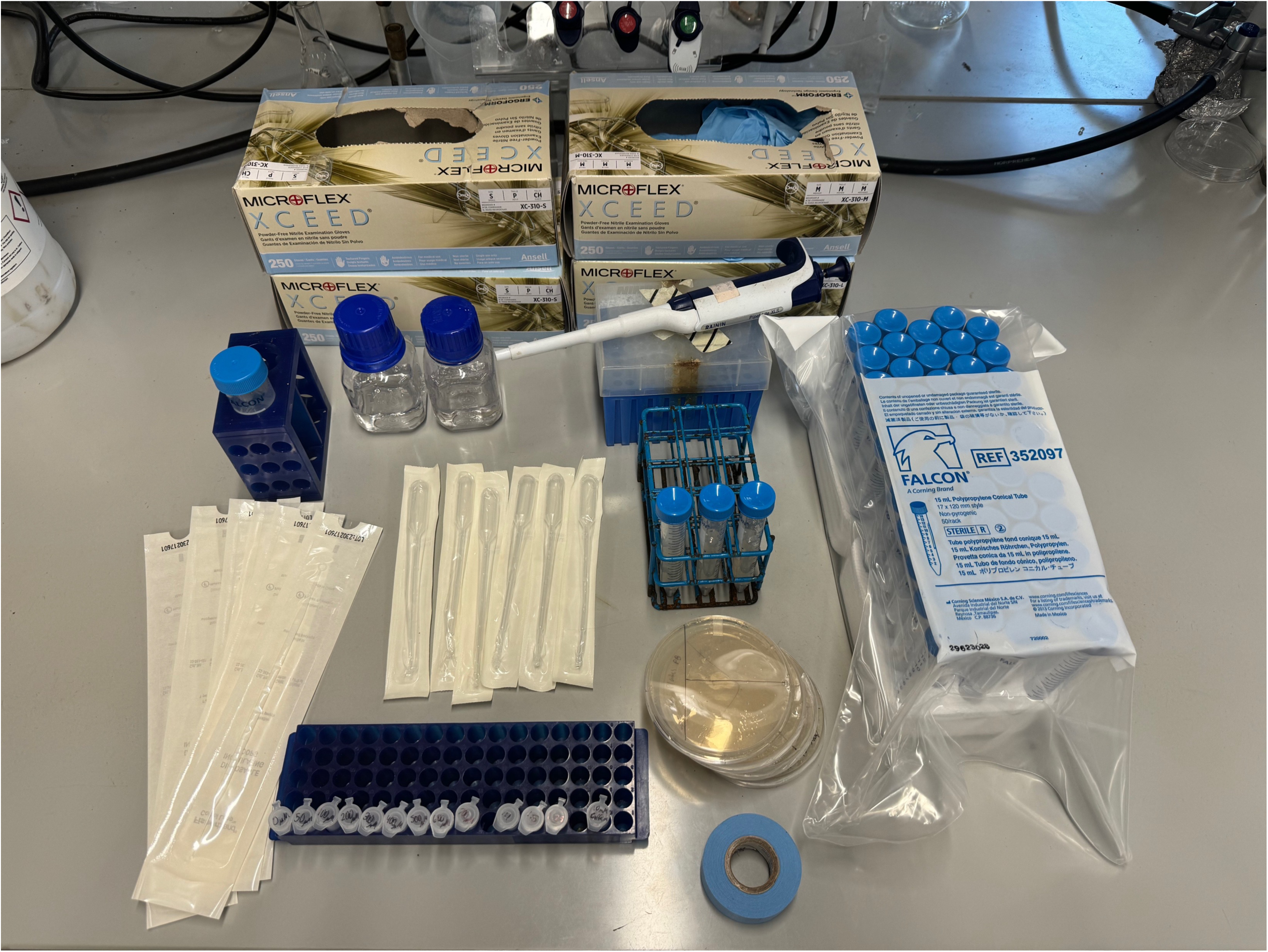
Caffeinated Coli Materials Needed for the Bioassay.

Before the bioassay begins, teachers prepare a starter culture by isolating bacteria from a pDCAF-ABC *E. coli* agar stab and growing the cells in media with caffeine at room temperature (20°-25°C) for around 24-72 hours, depending on the rate of bacterial growth. All teacher guides as well as detailed student lab procedures can be found online for free. Students then complete a pre-lab to familiarize themselves with the reagents and steps of the bioassay.

During the bioassay, students are divided into groups of four, with two members working on the standard curve and two on the calibration sample during week 3 (Figure 6). The standard curve team creates a set of cultures with known caffeine concentrations (0–70 μM) to establish a baseline for bacterial growth. The calibration sample team prepares cultures with unknown concentrations, which they will determine through data analysis using the standard curve’s linear trend line. The cultures are incubated at room temperature for 24-72 hours in a shaker incubator or can be hand-shaken hourly throughout the school day to simulate a shaker incubator. Students then measure bacterial growth using the ChromeBox they built or a spectrophotometer.

**Figure 6.**
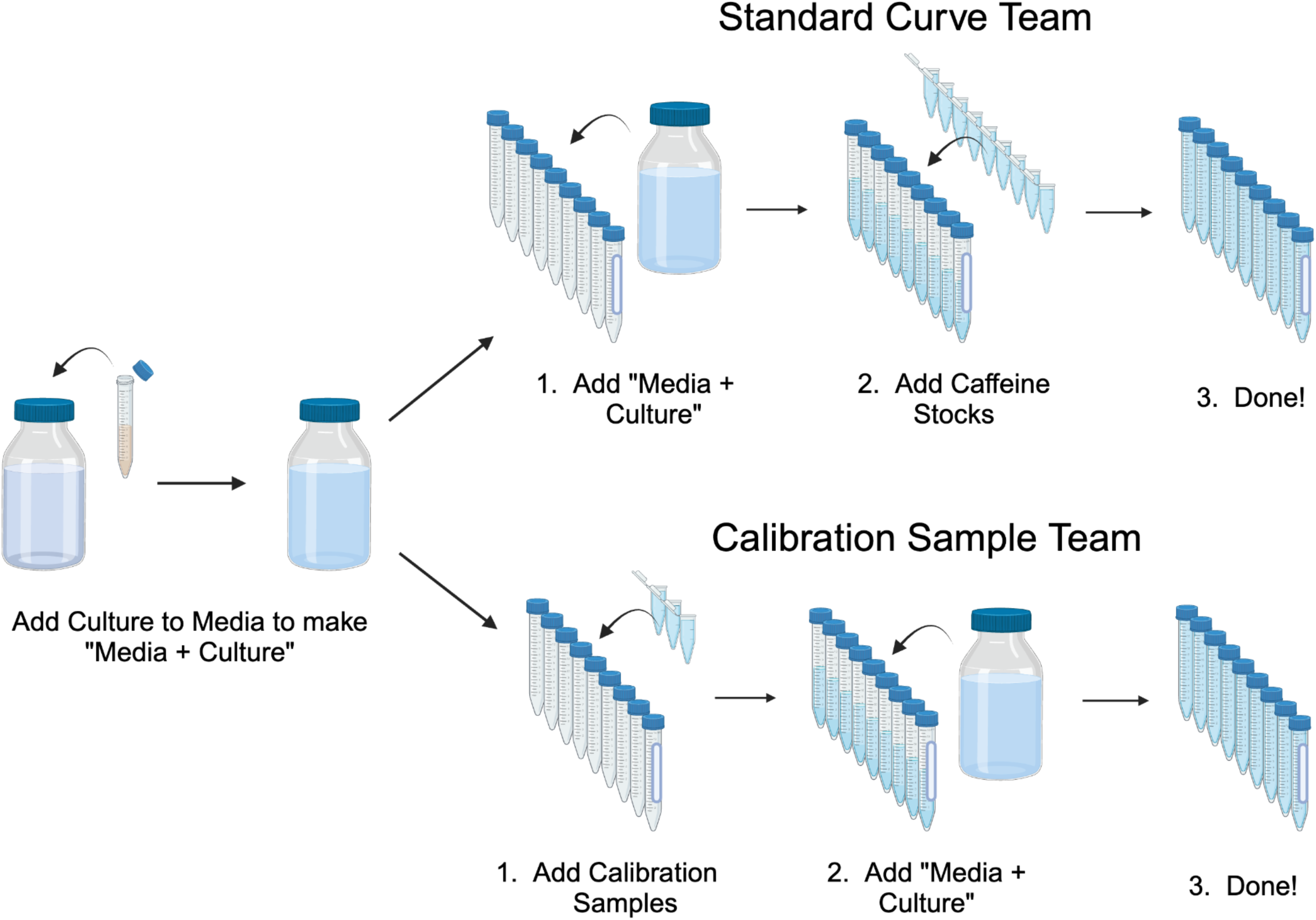
Caffeinated Coli Bioassay Pipeline. In Week 3, students create a standard curve and use pre-made calibration samples with a known concentration of caffeine.

In Week 4, students repeat the bioassay, this time using their own beverage samples instead of prepared calibration samples. The standard curve team creates a set of cultures with known caffeine concentrations, as they did in the first bioassay. The sample team (which replaces the calibration sample team from the first bioassay) then dilutes their beverages and adds the samples to culture tubes with media (Figure 7, bottom). Incubation again occurs for 24-72 hours, followed by data collection and analysis.

**Figure 7.**
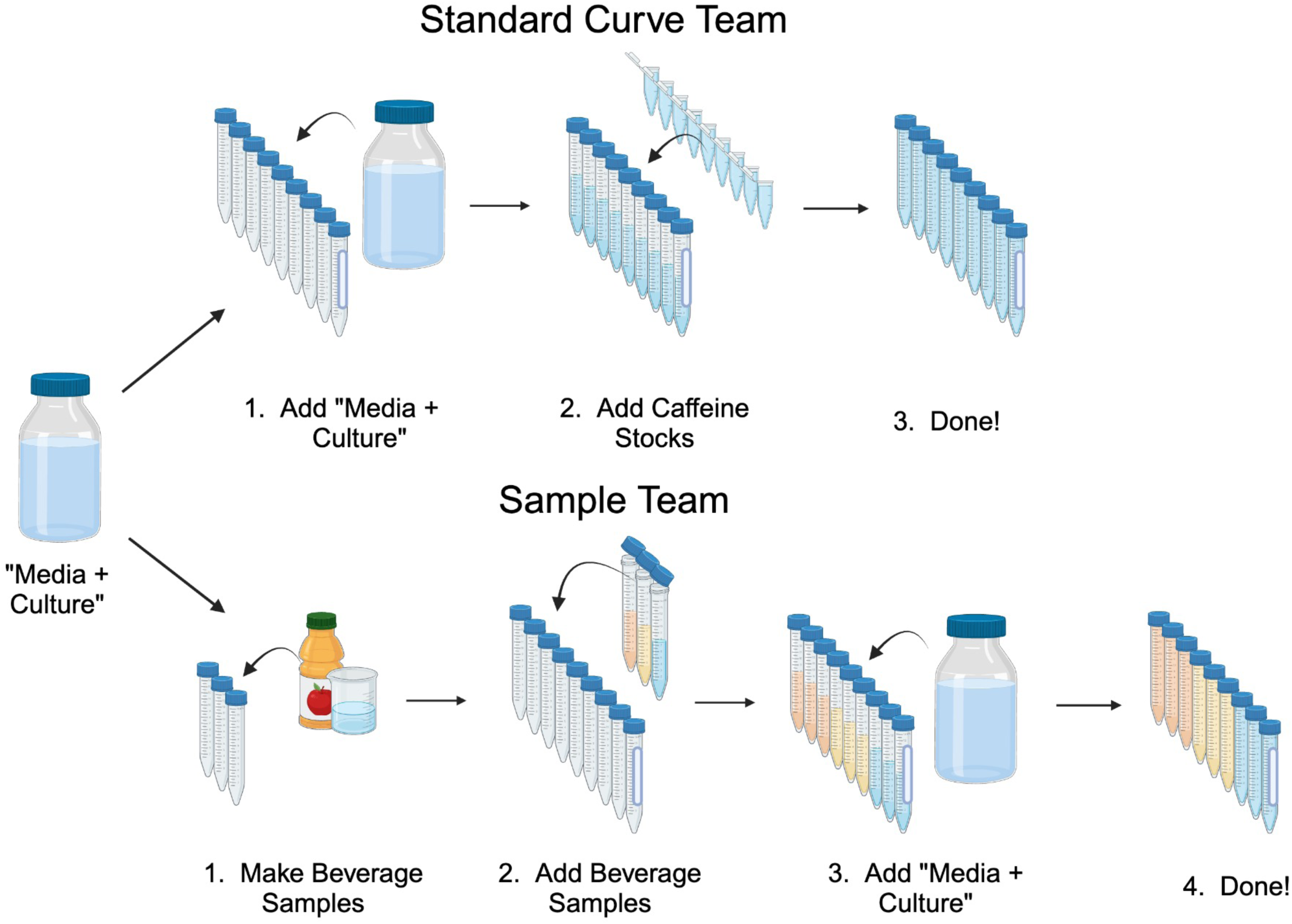
Week 4 Caffeinated Coli Pipeline. In Week 4, students create a standard curve and sample cultures using a beverage of their choosing.

All wet-lab procedures are conducted using sterile technique and with personal protective equipment (PPE), including lab coats, safety goggles, and gloves. All materials and strains needed for this module fall within standard BSL-1 safety guidelines, including biowaste disposal of the non-pathogenic *E. coli* being used. Suggested materials and reagents are not chemically hazardous, and no glassware, sharps, or needles are involved. As part of the module, we have included narrated, walk-through videos for the entire bioassay procedure to further ensure the safety and efficacy of the experiment. We have also eliminated the need for some potentially dangerous pieces of equipment such as a Bunsen burner, although we do have additional safety precautions, protocols, and demonstration videos if classes choose to incorporate Bunsen burners into their learning.

## Data Collection and Analysis (Week 4)

To measure the growth of each culture, students use a spectrophotometer or a lab-built ChromeBox, which is a cost-effective, interactive tool that students can build themselves using provided, simple instructions. It allows them to measure bacterial growth by using the “Colorimeter” smartphone app (see online resources, below), which reads color intensity through RGB values. Students work in groups to assemble the ChromeBox using basic materials, including a glove box, blue masking tape, and scissors. The device works by reflecting room light off the blue tape, which passes through their samples in a transparent tube (Figure 8). The Colorimeter app, positioned at the other end, measures the blue light passing through the sample (Soda, 2020).

**Figure 8.**
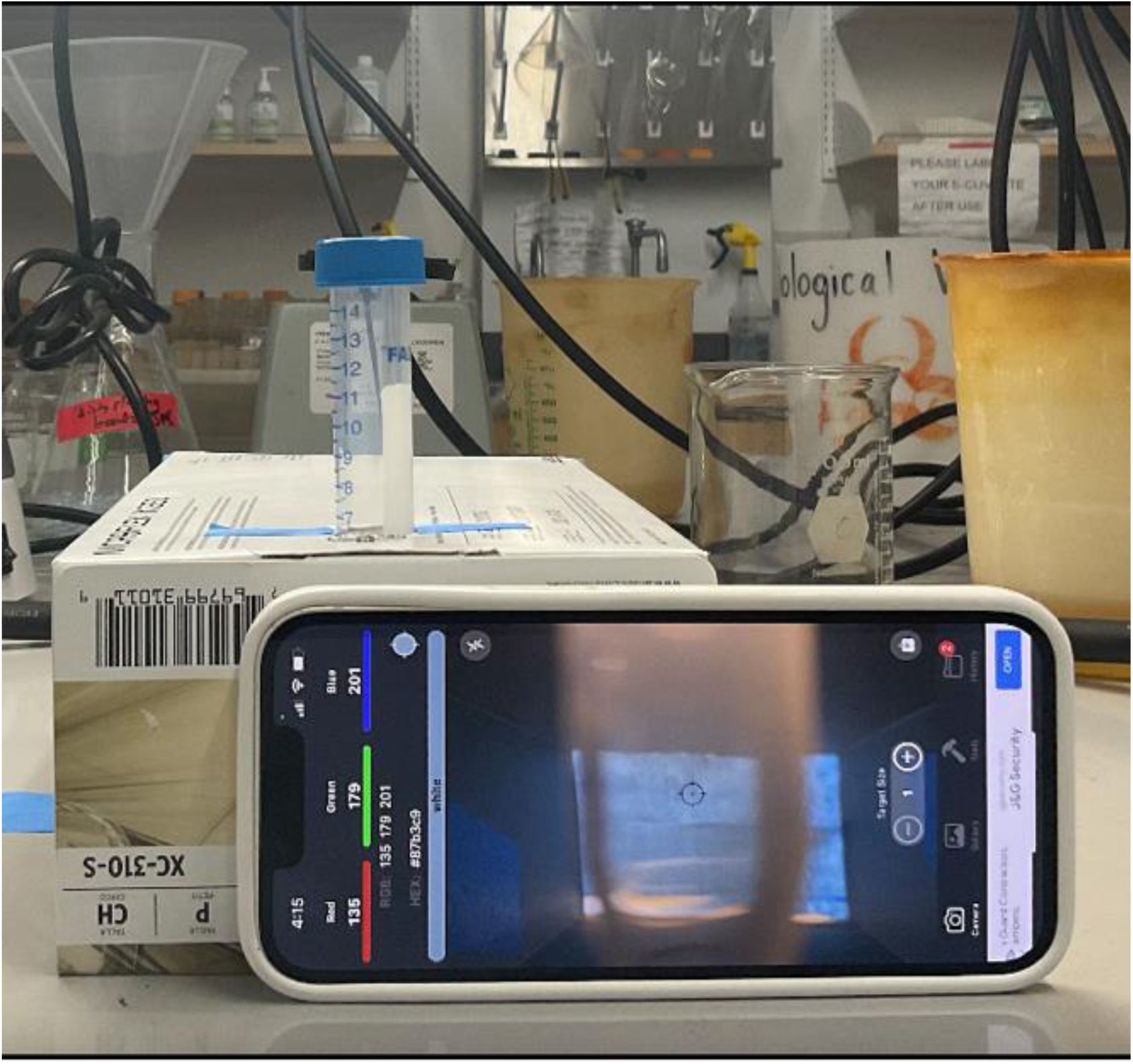
ChromeBox Display.

The amount of blue signal lost is directly proportional to the amount of cell growth in the culture, as more cells will block more light, similar to a spectrophotometer. After recording their readings in a spreadsheet, students follow a guided analysis, creating a standard curve and eventually determining the amount of caffeine in their sample.

## Scientific Communication (Week 5)

Students also learn how to present their findings through several activities. In the figure-making activity, they analyze examples of both strong and weak figures in a guided, class discussion, before creating their own figures and legends. Then, they share their figures with a partner, each assessing the quality of their partner’s figures, using guidelines to further their understanding of strong figures through the peer review process. Finally, students will work with their group members to create a final presentation, which not only allows students to showcase their experimental results but also allows them to communicate their knowledge and findings from the module, both of which are essential skills that are highly prized in research. Students are provided with an assignment outline, highlighting the expectations for their final presentation (Supplemental Table 3). We also provide the teacher with a rubric, to assist in grading. The rubric breaks down specific point values based on student data, figures, analysis, troubleshooting, and overall coherence (Supplemental Table 4). At the conclusion of the module, students engage in a brief reflection and receive resources related to how they can continue to pursue research in high school and beyond.

## Participant Feedback

Upon completing the Caffeinated Coli module, high school students and teachers evaluated its impact on their education and their overall experience through surveys. Students commonly reported the most memorable aspect of the module in 3 areas: engagement with mentors, collaboration in a group setting, and engagement with content/wet-lab work (Table 4). Teachers and students liked that students were given the opportunity to repeat the bioassay on two separate experimental days, including one where they chose the sample to assay. Office hours hosted by mentors – college students and a professor – were also beneficial as students were interested in asking questions regarding the college experience, medical school applications, and opportunities to conduct research in high school and beyond. In addition to live office hours, students highlighted that they enjoyed collaborating within a team setting to perform the bioassay and found it helpful to learn from their team members.

**Table 4.** HRI Survey Student Feedback.

| **Most Memorable Experience** | **Student Feedback** |
| --- | --- |
| **Engagement with Mentors** | “I loved the mentors, specifically their positive attitude and including all the students.”<br>“I enjoyed learning and gaining insight from current UT students who have participated in college-level labs and some research, as they shared the opportunities Freshman Research Initiative has offered them, which was inspiring to hear as a future Biology major.”<br>“They made me more interested because they allowed me to talk to older students and they offered advice and future career opportunities.” |
| **Collaboration in a Group Setting** | “One of the most memorable parts from The University of Texas experiments were from collaborating with classmates to complete the task.”<br>“I thought that each experiment had a good amount of collaboration, which helped further enhance the memorable parts of the experiments.” |
| **Engagement with content and wet-lab work** | “It was very visual and hands-on, and I enjoyed the connection between science and math through the graphs and derivatives to study our data.”<br>“I enjoyed the complexity of the lab with multiple parts and multiple disciplines including synthetic biology and statistics, which piqued my interest in such sub-specialties of science.” |

## Modularity and Flexibility in Curriculum

The 5-week module has been adapted to accommodate various class schedules and timings. Each lesson is designed to fit within a 50-minute class period, yet the curriculum can be adapted to block schedules or class periods outside the allotted period. Teachers who have implemented the curriculum have been able to easily combine lessons to fit larger class times. For instance, teachers have combined the data analysis lesson with the figure-making lesson within the same class period. Each lesson has overlap from topics covered the previous days, which provides students with a refresher and allows for a smoother transition between lessons. Additionally, multiple teachers have opted to only do some of the lessons from the first two weeks, focusing on the concepts that they have not yet covered with their students, while skipping concepts students have learned in previous years or earlier in the academic year. This flexibility has also allowed the module to be implemented in an array of courses at various grade levels (Table 5).

**Table 5.** Course and Grade Level Implementation.

| **Course** | **Grade level** |
| --- | --- |
| **Biology**<br>**(On level & Honors)** | 9th grade |
| **Chemistry** | 10th and 11th grade |
| **Medical Microbiology** | 11th and 12th grade |
| **Biotechnology** | 11th and 12th grade |
| **Scientific Research**<br>**and Design** | 12th grade |

## Conclusion

Synthetic biology is a growing scientific discipline with applications in many sectors of the world, including medicine and agriculture. The 5-week Caffeinated Coli Educational Module introduces synthetic biology to high school students and seeks to inspire them to pursue research from a young age. Students not only practice fundamental laboratory techniques such as micropipetting, growing bacterial cultures, and conducting a bioassay but also engage in lecture activities that reinforce key concepts. They analyze data from their own unique samples and create scientific figures, gaining hands-on experience in data visualization. Additionally, they develop essential personal skills by collaborating in group work, refining their communication abilities, and conducting independent inquiries with samples of their choosing.

### Online Resources section

Colorimeter App (App store needed): https://apps.apple.com/ee/app/colorimeter-app/id1542365656

Curriculum content: https://sites.utexas.edu/microbe-hackers/caffeine-kit/

## Acknowledgements

This work would not have been possible without the help of the entire Microbe Hackers Caffeinated Coli research group members (2020-2025). In particular, Tyler Le, Chloe Swanson, Peyton Gill, and Aalaysiah Morrison for their invaluable mentorship and assistance in the development of the Caffeinated Coli curriculum and reagents as well as Kyle Kamanu and Lisa Naranjo for their assistance in preparing reagents and communicating with high schools. We would like to thank the University of Texas at Austin’s Freshman Research Initiative and the College of Natural Sciences for funding and curriculum support. We would also like to extend our gratitude to the UT Austin UTeach program and Deanna Buckley and Gwen Stovall of the UT Austin’s High School Research Initiative for their expertise and assistance in developing the curriculum, deploying the module and surveys, IRB documentation, and giving feedback on the manuscript. Without their logistical support and experience working with high school curriculum, teachers, and students, this work would not have been possible. Figures 1-4, 6, and 7 were originally created using BioRender.com. This work was supported by the National Institute of General Medical Sciences, the National Institutes of Health under Award Number R25GM129171/R25GM142078. The content is solely the responsibility of the authors and does not necessarily represent the official views of the National Institutes of Health.

## Statement on human subjects/IRB approval

This work was reviewed by the University of Texas at Austin’s Institutional Review Board and was classified as exempt. IRB ID#2016020036-MOD03. Exemption was granted as we do not collect identifying information from participants. Teachers and UT near-peer mentors have provided written informed consent to participate in the study. This study has received a *Waiver of Documentation of Informed Consent* for high school students and an “opt out” consent (effectively a waiver of parental permission) for parents because all HRI activities in which the student participants conducted were typical of regular school days, such as participating in class, completing coursework, and completing assessments, and all the activities/materials were vetted by their teachers, as per usual. Parents were sent an Informational Sheet for Families describing the program and the study and provided a chance to opt their child out of data collection (e.g., student survey and pre/post-test).

