## Supplemental Table 1 for "Caffeinated Coli: Inspiring the Next Generation of Scientists through Synthetic Biology"

**Supplemental Table 1. Lesson Alignment with TEKS.**

| Lesson | Texas Essential Knowledge and Skills (TEKS) |
|---|---|
| **Week 1: Basics of Synthetic Biology** | C1(B) apply science practices related to specialized topics of study to plan and conduct investigations or use engineering practices to design solutions to problems |
| **Week 1: Introduction to Caffeinated Coli** | C3(B) communicate explanations or solutions individually and collaboratively in a variety of settings and formats as appropriate to the specialized topic of study |
| **Week 2: Introduction to ChromeBox** | C1(D) use tools appropriate to the specialized topic of study; C1(E) collect quantitative data using the International System of Units (SI); |
| **Week 3: Standard Curves** | C1(F) organize quantitative or qualitative data using representations appropriate to the specialized topic of study |
