## Supplemental Table 2 for "Caffeinated Coli: Inspiring the Next Generation of Scientists through Synthetic Biology"

### Supplemental Table 2. Learning Standards Connections.

| Performance Expectations: Caffeinated *Coli* | | |
|---|---|---|
| **Primary Performance Expectations (PEs):**<br>**HS-LS1-1:** Construct an explanation based on evidence for how the structure of DNA determines the structure of proteins, which carry out the essential functions of life through systems of specialized cells.<br>**HS-LS1-6:** Construct and revise an explanation based on evidence for how carbon, hydrogen, and oxygen from sugar molecules may combine with other elements to form amino acids and/or other large carbon-based molecules.<br>**HS-LS2-2:** Use mathematical representations to support and revise explanations based on evidence about factors affecting biodiversity and populations in ecosystems of different scales.<br>**HS-LS3-1:** Ask questions to clarify relationships about the role of DNA and chromosomes in coding the instructions for characteristic traits passed from parents to offspring. | | |
| **Science & Engineering Practices (SEPs)** | **Disciplinary Core Ideas (DCIs)** | **Crosscutting Concept (CCCs)** |
| **Engaging in Argument from Evidence:**<br>• Evaluate the evidence behind currently accepted explanations to determine the merits of arguments.<br>**Analyzing and Interpreting Data**<br>• Apply concepts of statistics and probability (including determining function fits to data, slope, intercept, and correlation coefficient for linear fits) to scientific and engineering questions and problems, using digital tools when feasible.<br>**Using Mathematics and Computational Thinking**<br>• Create or revise a simulation of a phenomenon, designed device, process, or system.<br>**Obtaining, Evaluating, and Communicating Information**<br>• Communicate scientific information (e.g., about phenomena and/or the process of development and the design and performance of a proposed process or system) in multiple formats (including orally, graphically, textually, and mathematically). | **LS1.A: Structures and Function:**<br>• Systems of specialized cells within organisms help them perform the essential functions of life.<br>• All cells contain genetic information in the form of DNA molecules. Genes are regions in the DNA that contain the instructions that code for the formation of proteins, which carry out most of the work of cells.<br>**LS2-A: Interdependent Relationships in Ecosystems**<br>• Ecosystems have carrying capacities, which are limits to the numbers of organisms and populations they can support. These limits result from such factors as the availability of living and nonliving resources and from such challenges as predation, competition, and disease. Organisms would have the capacity to produce populations of great size were it not for the fact that environments and resources are finite. This fundamental tension affects the abundance (number of individuals) of species in any given ecosystem.<br>**LS3.A: Inheritance of Traits**<br>• Each chromosome consists of a single very long DNA molecule, and each gene on the chromosome is a particular segment of that DNA. The instructions for forming species' characteristics are carried in DNA. All cells in an organism have the same genetic content, but the genes used (expressed) by the cell may be regulated in different ways. Not all DNA codes for a protein; some segments of DNA are involved in regulatory or structural functions, and some have no as-yet known function. | **Cause and Effect:**<br>• Empirical evidence is required to differentiate between cause and correlation and make claims about specific causes and effects.<br>**Systems and System Models**<br>• Models (e.g., physical, mathematical, computer models) can be used to simulate systems and interactions—including energy, matter, and information flows—within and between systems at different scales<br>**Stability and change**<br>• Much of science deals with constructing explanations of how things change and how they remain stable. |

NGSS Lead States. 2013. *Next Generation Science Standards: For States, By States.* Washington, DC: The National Academies Press.
