## Supplemental Table 3 for "Caffeinated Coli: Inspiring the Next Generation of Scientists through Synthetic Biology"

### Supplemental Table 3. Final Presentation Assessment.

| Final Presentation Rubric | Points Assigned | Points Earned |
|---|---|---|
| **Introduction/Background**<br>• Introduce the audience to the field and project: What are you trying to do? What is the bigger picture/importance of your work? | **15** | |
| **Results: Data and figures**<br>• This includes all your data collected including pictures, graphs, etc. You also need appropriate legends that explain what we are looking at.<br>• Some important figures to consider including pictures of your standard/sample cultures tubes (Parts 1 and 2), standard curve graphs, dilution calculation table, final caffeine concentration, etc. | **30** | |
| **Results: Analysis and Discussion**<br>• Interpretation of your results/data. What do your results mean? What have you concluded from your data? What is your key result/finding? This section should be written in bullet form or short paragraphs describing your results. | **15** | |
| **Potential Errors/Troubleshooting**<br>• Describe any potential errors that might have risen. Are these technique errors? Did you troubleshoot? How might have these errors affected your data? What will you do differently next time? | **15** | |
| **Grammar/Spelling**<br>• Is your text error-free (spelling, grammar, punctuation)? Is the information presented on the slides coherent to the audience? How visually appealing is your presentation? Does it present as a narrative rather than a list of experiments? | **10** | |
| **Presentation**<br>• Are you able to finish your well-rehearsed presentation in 10 minutes? Did you communicate your ideas clearly? Were all your group members given an equal speaking role? | **10** | |
| **References**<br>• What research articles or other sources did you use? Cite them appropriately at the end of your presentation. | **5** | |
| **TOTAL POINTS SCORED:___________** | | |
