## Supplemental Table 4 for "Caffeinated Coli: Inspiring the Next Generation of Scientists through Synthetic Biology"

### Supplemental Table 4. Final Presentation Rubric.

| Final Presentation Rubric | Points Assigned | Points Earned |
|---|---|---|
| **Introduction/Background**<br>• No mention of background or the project- **0 pts**<br>• The background and project are mentioned but are vague and unclear, providing no connection to the importance of the work- **5 pts**<br>• The project is addressed, but the bigger picture is only briefly mentioned- **10 pts**<br>• The project goal is stated clearly, and its bigger picture is explained thoroughly- **15 pts** | **15** | |
| **Results: Data and figures**<br>• No data or figures are present- **0 pts**<br>• Some data is present but is incomplete, disorganized, or lacks legends- **10 pts**<br>• Most required data and figures are present but may be missing one or two key elements, or legends are minimal and lack clarity- **20 pts**<br>• All required data and figures are present with clear, complete legends that articulate what is shown- **30 pts** | **30** | |
| **Results: Analysis and Discussion**<br>• No analysis or discussion is present- **0 pts**<br>• Briefly interprets data, but the conclusions are unclear, illogical, or contradict the results presented- **5 pts**<br>• Results are summarized and interpreted, but the discussion lacks depth and fails to highlight key findings or misses connection to significance- **10 pts**<br>• Results are thoughtfully interpreted, key findings are fully supported by data, and highlights connection to the bigger picture- **15 pts** | **15** | |
| **Potential Errors/Troubleshooting**<br>• No potential errors, troubleshooting, or improvements are discussed- **0 pts**<br>• Errors are mentioned but are vague, lack a specific cause (technique, etc.), and do not explain their impact on the data- **5 pts**<br>• Errors are mentioned with a brief explanation on how they affect results, but troubleshooting and future improvements are not discussed- **10 pts**<br>• Specific errors, including their effect on data, troubleshooting methods, and improvements for future work are thoroughly described- **15 pts** | **15** | |
| **Grammar/Spelling**<br>• Text contains significant spelling/grammar errors and is visually disorganized- **0 pts**<br>• Text has several grammatical errors, and presentation lacks visual appeal or structure- **3 pts**<br>• The text has a few grammatical errors and information is clear, but the presentation lacks narrative- **7 pts**<br>• Text is error-free, visually appealing, and presents a clear, concise narrative- **10 pts** | **10** | |
| **Presentation**<br>• Presentation was unrehearsed and significantly over/under time- **2 pts**<br>• Presentation was rehearsed and within time, but unclear in a few parts or speaking role was unequal- **5 pts**<br>• Presentation was well-rehearsed, finished within time, and displayed clear communication with all members participating equally- **10 pts** | **10** | |
| **References**<br>• No references present or mentioned- **0 pts**<br>• At least one reference mentioned- **1 pts**<br>• References mentioned or listed in most places, as appropriate- **3 pts**<br>• References appropriately mentioned or listed throughout presentation- **5 pts** | **5** | |
